# Steroidal glycoalkaloid biosynthesis and fungal tolerance are regulated by ELONGATED HYPOCOTYL 5, SlHY5, in tomato

**DOI:** 10.1101/2024.02.29.582793

**Authors:** Hiteshwari Sinha, Ravi Shankar Kumar, Tapasya Datta, Deeksha Singh, Suchi Srivastava, Prabodh Kumar Trivedi

## Abstract

Tomato (*Solanum lycopersicum* L.) is one of the highest consumable fruit crops, rich in nutrients, and has been an important target for enhancing the accumulation of various metabolites. Tomato also contains cholesterol-derived molecules, steroidal glycoalkaloids (SGAs), which contribute to pathogen defence but are toxic to humans and considered anti-nutritional compounds. Previous studies suggest the role of various transcription factors in SGA biosynthesis; however, the role of light and associated regulatory factors has not been studied in tomatoes. Here, we demonstrated that SGA biosynthesis is regulated by light through the ELONGATED HYPOCOTYL 5 homolog, SlHY5, by binding to light-responsive G-boxes present in the promoters of the structural and regulatory genes. Our analysis suggests that SlHY5 could complement the *Arabidopsis thaliana* and *Nicotiana tabacum, hy5* mutants at molecular, morphological, and biochemical levels. We report the development of CRISPR/Cas9-based knockout mutant plants of tomato, *slhy5^CR^*, and show down-regulation of the SGA and phenylpropanoid pathway genes leading to a significant reduction in SGA (α-tomatine and dehydrotomatine) and flavonol contents, whereas SlHY5 overexpression (SlHY5OX) plants show opposite effect. An enhanced SGA and flavonol levels in SlHY5OX lines provided tolerance against *Alternaria solani* fungus, while *SlHY5^CR^* was susceptible to the pathogen. This study advances our understanding of the HY5-dependent light-regulated biosynthesis of SGAs and flavonoids and their role in biotic stress in tomatoes.

**One Sentence Summary:** Light-associated transcription factor, ELONGATED HYPOCOTYL 5, regulates biosynthesis of anti-nutrient molecules, steroidal glycoalkaloids, and fungal tolerance in tomato

## INTRODUCTION

Secondary metabolites are synthesized and accumulated in plants that predominately function to shield plants from internal and external injuries, which may be biotic or abiotic in nature (Winkel-Shirley, 2002; Tanaka et al., 2008; Misra et al., 2010; Sharma et al., 2023). Alkaloids are one of the major classes of secondary metabolites produced in the plant. Among alkaloids, Steroidal Glycoalkaloids (SGAs), mostly found in the Solanaceae family, are classified by their nitrogenous steroidal aglycone and glycoside residues located at the C-3 position. The SGAs are well known for their antinutritional properties for humans and animals. However, they can also be used as pharmaceutical compounds because of their diverse biological roles, for instance, anti-inflammatory, anti-microbial, anti-fungal, anti-cholesterol, cytotoxic, inhibitor of skeletal muscle atrophy, and anti-pyretic effects (Milner et al., 2011; Friedman 2006; Ito et al., 2007; Bailly 2021). To date, more than a hundred SGAs have been identified in Solanum species, and in tomatoes, they accumulate in various plant organs such as leaves, flowers, stems, roots, and green fruits. Dehydrotomatine and α-tomatine are the two SGAs that are primarily present in tomatoes (Zhao et al., 2021; Sonawane et al., 2020; Yu et al., 2020; Wang et al., 2018; Cardenas et al., 2016).

Cholesterol, the main precursor for the biosynthesis of SGAs, is synthesized using acetyl-CoA as the carbon source. Through various enzymatic reactions of hydroxylation, oxidation, and transamination, cholesterol forms the steroidal aglycone containing glycoside residues catalyzed by the specific gene cluster family, Glycoalkaloid Metabolism (GAME) **(Figure 1A)** (Itkin et al., 2013; Sawai et al., 2014; Wang et al., 2018). In the Solanaceae family, GAME9 (Jasmonic acid-responsive ERF4) plays a crucial regulatory role in the SGA biosynthesis as the loss of function of GAME9 resulted in a considerable drop in alkaloid production (Cardenas et al., 2016; Thagun et al., 2016; Nakayasu et al., 2018). Previous reports suggested that potato tubers exposed to light accumulate more glycoalkaloid compared to control plants, suggesting an important role of light in SGA biosynthesis. In plants, light exposure induces the transcriptional reprogramming of approximately 20-35% of genes through the interaction of various transcription factors to LREs (light-responsive elements) present in the promoter of light-responsive genes (Quail, 2002; Leivar and Quail, 2011; Gangappa et al., 2013a; Gangappa and Botto, 2014).

**Figure 1.**
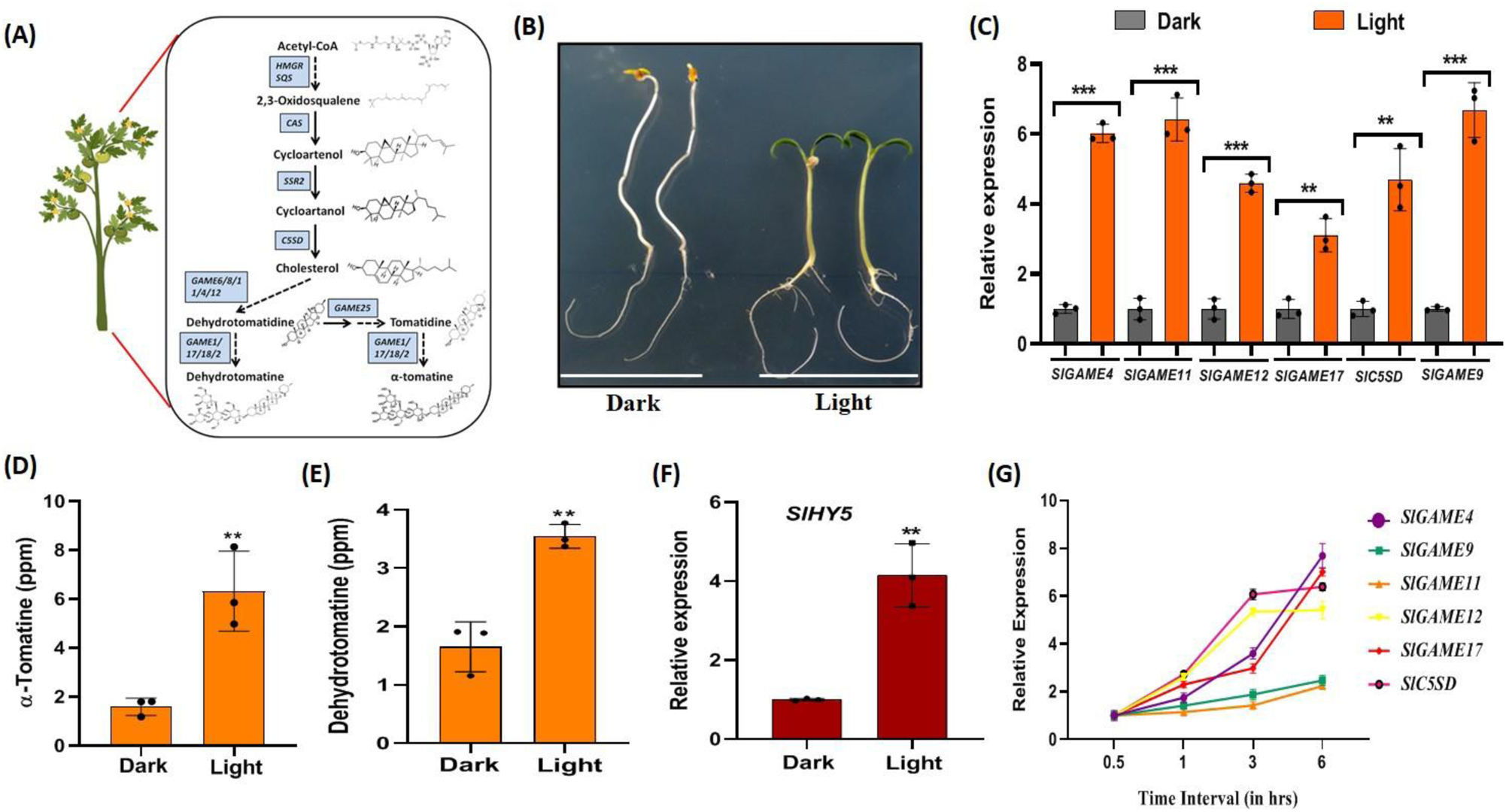
Light-mediated regulation of SGA biosynthesis in WT tomato. **A,** Pictorial representation of SGA biosynthesis pathway of tomato. **B,** Phenotype of dark and light grown WT seedlings after 10 days. **C,** Expression analysis of SGA pathway structural and regulatory genes (*SlGAME4, SlGAME11, SlGAME12, SlGAME17, SlC5SD* and *SlGAME9*) in 10-days-old light-and dark-grown WT seedlings. **D-E,** Quantitative estimation of α-tomatine and dehydrotomatine in 10-day-dark and light-grown WT seedlings through LC-MS. **F**, Relative expression of *SlHY5* in 10-d-dark and light grown seedlings. **G,** Time dependent qRT-PCR expression analysis of SGA pathway genes (*SlGAME4, SlGAME11, SlGAME12, SlGAME17, SlC5SD* and *SlGAME9*) in 10-day dark grown WT tomato seedlings consequently transferred to light for 0.5, 1, 3, and 6 hours. Statistical analysis was performed using two-tailed Student’s t-test. Error bars represent SE of means (n=3). Actin was used as endogenous control to normalize the relative expression levels. Error bars represent standard deviation. Asterisks indicate a significant difference, *P < 0.1, **P < 0.01, ***P < 0.001.

In recent times, one of the key light-regulated transcription factors (TF) is known as HY5 (Elongated Hypocotyl 5), belonging to the bZIP transcription factor family, has been extensively studied as it plays a pivotal role in plant growth and development via regulating the light-responsive transcriptional reprogramming (Xiao et al., 2022; Jiao et al., 2007; Heijde and Ulm., 2012; Lau and Deng., 2012; Gangappa and Botto, 2014). HY5 positively regulates various biological processes such as photomorphogenesis, lateral root growth development, and root gravitropism. In *Arabidopsis*, HY5 acts as a master regulator by binding to cis-regulatory elements of more than 3000 genes directly (Zhang et al., 2011) and regulates flavonoid and terpenoid biosynthesis (Bhatia et al., 2018; Michael et al., 2020). The stability of HY5 is regulated by E3 ubiquitin ligase, COP1 (CONSTITUTIVE PHOTOMOROHOGENIC1), that leads to ubiquitination and 26S proteasomal degradation of HY5 in the dark although, in the ambient light condition, COP1 is negatively regulated by HY5 which promotes photomorphogenic response. Thus they act as an antagonist in plants (Chen et al., 2013; Xu et al., 2014; Hoecker, 2017; Bhatia et al., 2021). The extensive characterization of HY5 and secondary plant products have been done in *Arabidopsis*, rice, pea, potato, eggplants, tobacco, and lotus (Burman et al., 2018; Yamawaki et al., 2011, Nishimura et al., 2002; Weller et al., 2009; Zhai et al., 2020; Singh et al., 2022) but, the characterization of tomato HY5 and its role in light-regulated photomorphogenesis as well as in SGAs/flavonoid biosynthesis require further insight.

In this study, to decipher the function of HY5 of tomato in SGA and flavonoid biosynthesis, we identified and functionally characterized SlHY5 through complementing *Arabidopsis* and tobacco *hy5* mutants (*athy5* and *nthy5,* respectively), developing SlHY5-overexpressing (SlHY5OX) lines and CRISPR/Cas9-based mutants (*slhy5^CR^*). CRISPR/Cas9-based HY5 mutants accumulated significantly lower amounts of SGAs compared to wild-type tomato plants, suggesting the involvement of light-associated signaling in the regulation of SGA biosynthesis. We demonstrate that SlHY5 acts as a major light-responsive factor that regulates the light-mediated SGA/flavonoid biosynthesis in tomato by interacting with promoter regions of genes involved in SGA/flavonoid biosynthesis. As SGAs/flavonoids are known to provide biotic stress tolerance, we show that changes in levels of these metabolites in SlHY5OX and *slhy5^CR^*plants lead to modulated response towards *Alternaria solani,* that may lead to the most devastating early blight foliar disease causing defoliation and reduced yields in tomato (Jones et al., 2014; Jones and Perez, 2019; Attia et al., 2020).

## RESULTS

### Light-dependent SGA and flavonoid biosynthesis in tomato

To investigate the influence of light on SGA biosynthesis in tomato, the WT tomato seedlings were grown in light and dark for 10 days, which showed significant phenotypic variation in their hypocotyls, chlorophyll, and cotyledons **(Figure 1B).** The expression analysis of the genes involved in the SGA biosynthesis showed enhanced expression of structural (*SlGAME4/11/12/17* and *SlC5SD*) and, regulatory genes (*SlGAME9*) **(Figure 1C)** in light-grown compared to dark-grown seedlings. Similarly, an enhanced expression was observed in the structural and regulatory genes involved in the flavonoid biosynthesis (*SlCHS, SlCHI, SlFLS, SlF3H, SlDFR* and *SlMYB12*) **(Supplemental Figure S1A)** in light-grown seedlings compared to dark-grown seedlings. Quantification of SGAs content (α-tomatine and dehydrotomatine) in the light and dark-grown seedlings showed a significant increase of α-tomatine and dehydrotomatine content in seedlings grown in light compared to the dark **(Figure 1D-E).** The expression of *SlHY5*, a major light-associated regulatory factor, was much higher in light-grown compared the dark-grown seedlings **(Figure 1F).** Since flavonol accumulation is known to be regulated by light (Pandey et al., 2014; Bhatia et al., 2018), flavonol content (kaempferol and quercetin) was measured in the seedlings. A higher accumulation of flavonols was observed in light-grown compared to dark-grown seedlings **(Supplemental Figure S1B).**

To study the effect of light on the biosynthesis of SGA and flavonoid biosynthesis genes, the 10-day-old dark grown seedlings was exposed to light for different time intervals. The analysis suggested exposure of light to dark-grown seedlings enhances the expression of genes of SGA **(Figure 1G)** and flavonoid pathway genes **(Supplemental Figure S1C)**. We also analysed the expression of flavonoid biosynthetic genes and flavonol content in 10-day-old light-grown seedlings of the AtMYB12OX line of tomato (Pandey et al., 2015) exposed to the dark for 24 and 48 h. Analysis suggested a decreased expression of flavonoid biosynthesis genes and flavonol content in dark-exposed compared to light-grown seedlings, suggesting the involvement of light in secondary metabolite production in addition to MYB transcription factor **(Supplemental Figure S2 and S3).**

Since HY5 is well known as a key regulator of light-associated responses in numerous plants, we identified the tomato HY5 (SlHY5) to analyse its involvement in SGA and flavonoid biosynthesis. Sequence analysis suggested that SlHY5 has more than 90% homology with *Arabidopsis* and tobacco HY5 (AtHY5 and NtHY5) and contains conserved leucine-rich zinc finger and VPE motifs (Holm et al., 2002), indicating that it belongs to the bZIP superfamily like other plants and interact with COP1 **(Supplemental Figure S4).** Phylogenetic and pairwise distance analysis using different plant species suggested that SlHY5 is closer to NtHY5 than AtHY5. Additionally, SlHY5 shared a close resemblance with HaHY5 and CtHY5 **(Supplemental Figure S5A-B).**

### SlHY5 is a functional ortholog of AtHY5 and NtHY5

Since SlHY5 showed sequence similarity and shared conserved domains with HY5 of different plant species, we hypothesized that the complementation of SlHY5 in the *hy5* mutants might complement associated functions and phenotype. To explore the biological function of SlHY5, we generated complemented lines of *Arabidopsis hy5* mutant (*athy5* background) and tobacco *hy5* mutant (*nthy5;* Singh et al., 2022) and were analyzed thoroughly for gene expression, biochemical aspects and phenotypic changes **(Supplemental Figure S6A).** The hypocotyl length of 10-day-old *Arabidopsis* and tobacco got restored to WT in complemented lines **(Supplemental Figure S6B).** The *Arabidopsis* hypocotyl length was nearly 0.4, 1.2, and 0.5 cm, and in tobacco, the average lengths were 0.2, 0.5, and 0.2 cm for WT, *hy5*, SlHY5:*hy5* seedlings, respectively. According to Zhang et al., (2017), AtHY5 is known for light-dependent root growth development; hence the root length of 10-day-old seedlings was evaluated in these lines of *Arabidopsis* and tobacco, suggesting the recovered root lengths of complemented line to WT seedlings **(Supplemental Figure S6C).** Rosette diameter (4, 7, 5 cm for WT, *hy5*, SlHY5:*hy5* of *Arabidopsis,* respectively) **(Supplemental Figure S7)** and plant height of mature plant suggested recovery of the lost phenotype **(Supplemental Figure S8).** Correspondingly, a similar result was observed with seed size **(Supplemental Figure S9A-B**). All these data showed that SlHY5 complements the phenotypes of AtHY5 and NtHY5 in their *hy5* mutant.

To validate the SlHY5 function at the transcript and biochemical aspects, we examined the expression of genes involved in the phenylpropanoid pathway in 10-day-old seedlings of WT, *hy5*, and SlHY5:*hy5* complemented lines of *Arabidopsis* and tobacco. Based on earlier reports, mostly common phenylpropanoid are synthesized in different plants; however, synthesis of a few plant-specific metabolites have also been reported (Tohge et al., 2017; Xiao et al., 2022) (**Supplemental Figure S10**). The expression analysis revealed that the expression of flavonoid biosynthesis genes (*AtCHS/NtCHS, AtCHI/NtCHI, AtFLS/NtFLS,* and *AtMYB12/NtMYB12*) re-established as the WT in complemented lines of *Arabidopsis* and tobacco **(Supplemental Figure S11A-B)** and the accumulation of kaempferol and quercetin regained as WT in *Arabidopsis* and tobacco **(Supplemental Figure S12A-B; Supplemental Figure S13A-B).** While in tobacco, similar results were observed for other flavonols (rutin and CGA) **(Supplemental Figure S12C).** Since HY5 is related to light signaling, seeds were grown in the dark for 10 days and seedlings were transferred in light for 6 hours. The expression of flavonoid biosynthesis genes (*AtCHS* and *AtFLS*) were significantly upregulated on light illumination in complemented lines; however, on the contrary, the *hy5* mutant did not show a significant increase of these genes at the transcript level **(Supplemental Figure S14A).** Besides, the level of the transcript of the key transcription factor of the MYB family, *AtMYB12*, also increased as WT in complemented lines **(Supplemental Figure S14B)**. Since previous reports suggest that the chlorophyll and anthocyanin biosynthesis is regulated by AtHY5 (Oyama et al., 1997), we also measured the total flavonoid and chlorophyll content in 10-day-old light-grown seedlings of WT, *hy5* mutant, and complemented lines. The total chlorophyll and flavonoid content in complemented lines was greater than the *hy5* mutant and similar to the WT **(Supplemental Figure S15A and B)**. These results indicated that SlHY5 could recover for the loss not only at transcript level but also at metabolite level too in *athy5* and *nthy5* mutants and complements the function of HY5 in plants.

### Development and analysis of *SlHY5* overexpression and SlHY5-edited lines

To further investigate the functions of SlHY5 and its involvement in SGA and flavonoid biosynthesis, *SlHY5* mutant plants (*SlHY5^CR^*) using the CRISPR/Cas9 approach and *SlHY5* overexpression (SlHY5OX) plants were generated. To generate mutation in SlHY5, we designed guide RNAs (gRNAs) targeting the coding region of SlHY5 **(Supplemental Figure S16A)**. The nucleotide sequence analysis of the mutant plants revealed deletions and substitutions in the coding regions of SlHY5 **(Supplemental Figure S16B).** The edited lines of *SlHY5* plants showed frame-shift mutations leading to truncated peptides **(Supplemental Figure S16C)**. To develop overexpression lines of *SlHY5* in tomato, we cloned the coding region of SlHY5 under the CaMV35S promoter **(Supplemental Figure S17A).** All the lines were grown till T3 generation and used for further analysis. To analyze the effect of SlHY5 on plant growth and development, the WT, SlHY5OX, and *SlHY5^CR^* grown in half-strength MS medium and their phenotypic study were performed. The mutants showed longer hypocotyls, whereas, SlHY5OX represented shorter hypocotyls compared to the WT **(Figure 2A).** Upon maturation, the SlHY5OX showed reduced height and *SlHY5^CR^*plants represented increased height compared to the WT, indicating that SlHY5 regulates tomato plant growth and development **(Supplemental Figure S17B).**

**Figure 2.**
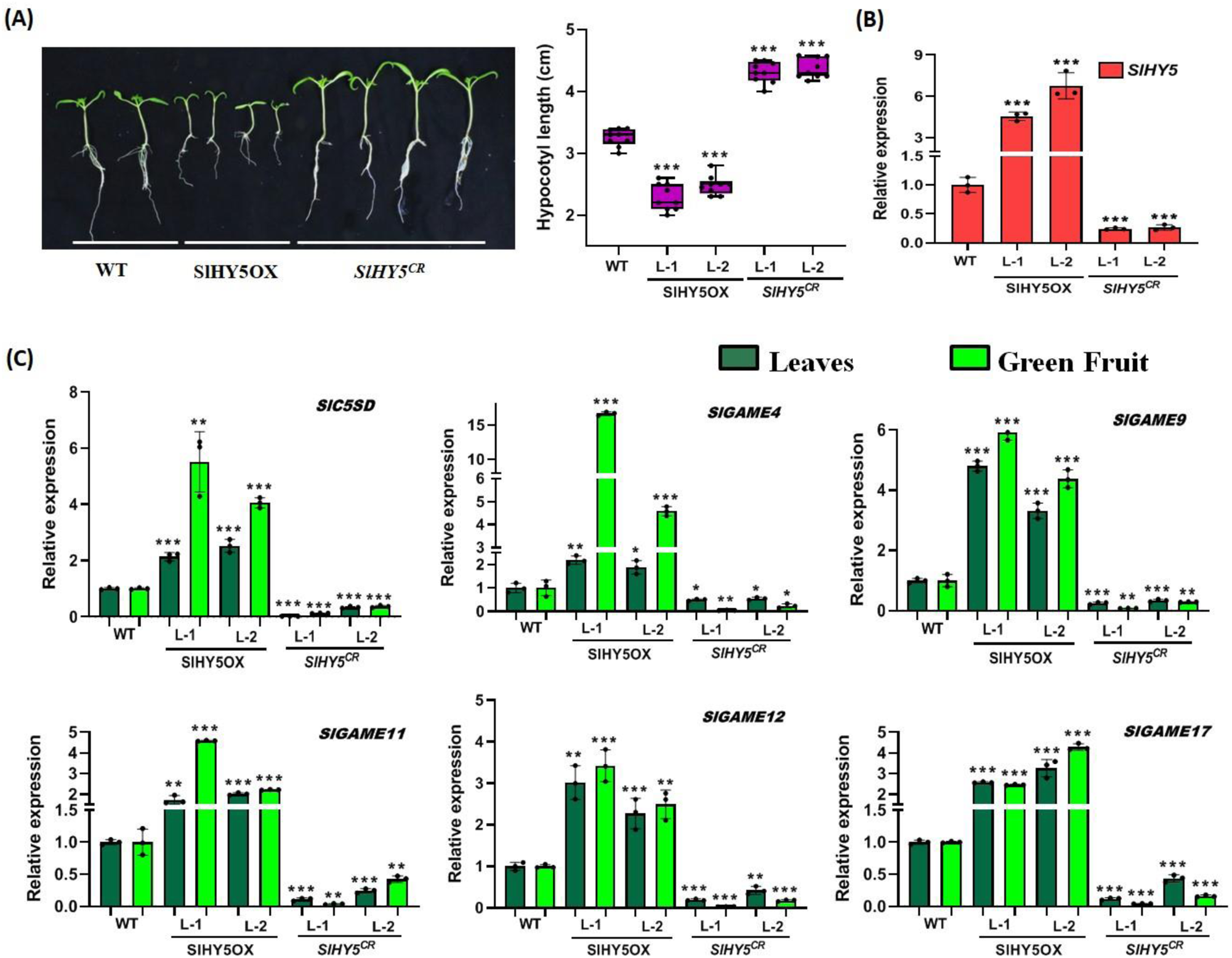
Overexpression of *SlHY5* and CRISPR/Cas9-mediated knockout of *SlHY5* modulates SGA pathway gene expression in tomato. **A,** phenotypic and graphical representation of hypocotyls length of the 10-days-old seedlings of tomato. **B,** Relative expression of *SlHY5* in 10-day-old light grown WT, SlHY5OX and *SlHY5^CR^*lines of tomato. **C,** Relative transcript abundance of SGA pathway genes (*SlGAME4, SlGAME11, SlGAME12, SlGAME17, SlC5SD* and *SlGAME9*) in 1-month-old light grown WT, SlHY5OX and *SlHY5^CR^*lines (leaves and green fruit). Actin was used as the endogenous control to normalize the relative expression levels. The statistical analysis was performed using two-tailed Student’s t-tests. The data are plotted as means ± SD (n=3). The error bars represent standard deviations. The asterisks indicate significant difference, *P< 0.1; **P< 0.01; ***P< 0.001.

The relative expression of *SlHY5* in SlHY5OX and *SlHY5^CR^*plants showed higher and lower transcript levels compared to the WT, respectively **(Figure 2B).** Additionally, to study the SlHY5 accumulation, the protein content in 10-days-old seedlings of SlHY5OX and edited lines was analysed using immunoblot assay. The result showed that overexpression of *SlHY5* led to significantly enhanced HY5 content whereas, edited lines showed no HY5 protein accumulation **(Supplemental Figure S18A)**. Based on the relative transcript levels, one-month-old young leaves and green fruits were used to analyse the expression of genes involved in SGA biosynthesis. The analysis implied that the expression of the gene cluster family of glycoalkaloid metabolism (*GAME4/9/11/12/17* and *SlC5SD*) was significantly lower in the *SlHY5^CR^*while SlHY5OX lines showed higher expression in young leaves and fruits compared to the WT **(Figure 2C)**. Moreover, flavonoid pathway structural genes like *SlCHS, SlCHI, SlFLS,* and regulatory gene, *SlMYB12*, were decreased in *SlHY5^CR^* seedlings, whereas a contrasting result was observed in overexpression lines **(Supplemental Figure S18B-D).** Biochemical analysis of 10-day-old seedlings showed that SlHY5OX lines accumulated higher chlorophyll, flavonoid content, and enhanced antioxidant activity, whereas the *SlHY5^CR^* lines show lower accumulation of chlorophyll, flavonoids and decreased antioxidant activity compared to the WT **(Supplemental Figure S19A-C).**

### SlHY5 interacts with the promoters of SGA and flavonoid biosynthesis genes

Our results suggested that the expression of SGA and flavonoid biosynthesis genes is higher in light-grown and SlHY5OX compared to dark-grown and *SlHY5^CR^* seedlings. Consequently, we were inquisitive to know whether the light regulator, SlHY5, directly regulates these genes. Our *in*-*silico* analysis suggested the presence of potential light-responsive elements (GATA, Box 4, ACE, and GT1) **(Supplemental Table S1-S7)** in the promoters of SGA and flavonoid biosynthesis genes. Further analysis suggested the presence of G-boxes (ACGT) in the promoters of *SlGAME11, SlGAME17, SlGAME4, SlGAME12, SlC5SD* and the major regulator of SGA biosynthesis, *SlGAME9* **(Figure 3A)**. Flavonoid biosynthesis regulator, *SlMYB12*, comprises one G-box at −825 bp **(Supplemental Figure S21A).**

**Figure 3.**
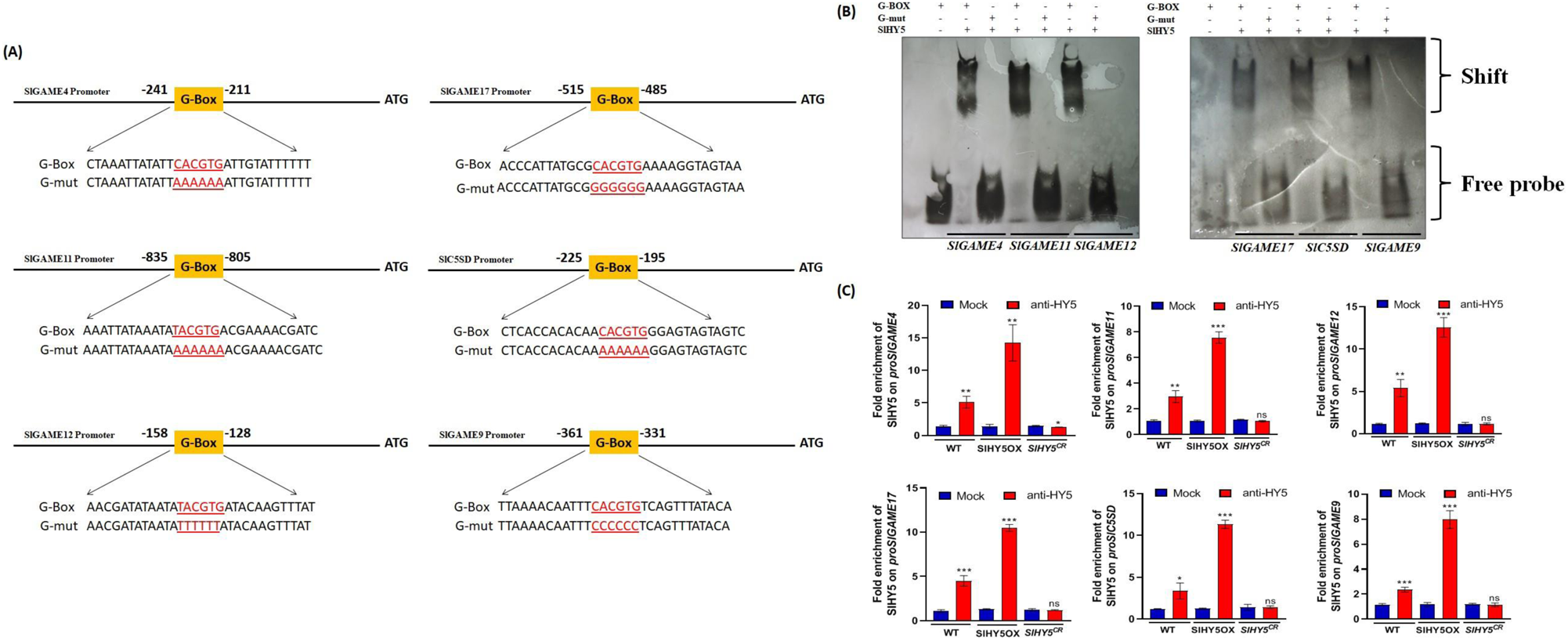
*In vitro* interaction between SlHY5 and the ACGT motif of G-BOX present at the promoters of SGA pathway genes. **A,** Representation of the probes (G-BOX, G-mut) designed upstream of the transcription start site of *SlGAME4*, *SlGAME11*, *SlGAME12*, *SlGAME17*, *SlC5SD* and *SlGAME9*. G-BOX is altered by multiple base substitutions. **B,** EMSA (Electrophoretic mobility shift assay) for the binding of 6X-His-HY5 (SlHY5) with core ACE DIG element (HY5-binding site that is G-BOX) present in the LRE motifs of the *SlGAME4*, *SlGAME11*, *SlGAME12*, *SlGAME17*, *SlC5SD* and *SlGAME9*, respectively. Upper and lower bracket indicate shift and free probe respectively. **C,** Fold enrichment of SlHY5 on promoters of SGA biosynthesis genes using ChIP-qPCR.

To validate the interaction, 6XHis-tagged SlHY5 was expressed in *E. coli* and purified protein **(Supplemental Figure S20A-C)** was used for electrophoretic mobility shift assay (EMSA). EMSA assay revealed that DIG-labeled G-box probes (G-BOX) for all SGA biosynthesis pathway genes, as well as *SlMYB12*, bind to *SlHY5* protein **(Figure 3B; Supplemental Figure S21B)**. At the same time, no band shift was observed with the mutated probes. Additionally, upon increasing the concentration of the cold probe (unlabeled probe) of *SlGAME4/9/11/12/17, SlC5SD* **(Supplemental Figure 22A-F)**, and *SlMYB12* **(Supplemental Figure S21C)**, the intensity of binding got diminished.

Additionally, we employed ChIP-qPCR to investigate the recruitment of SlHY5 to the promoters of SGA and flavonoid biosynthesis genes. Our analysis indicates that SlHY5 is indeed recruited to these promoters. Notably, the recruitment is significantly higher in SlHY5OX lines compared to WT plants, while *SlHY5^CR^* plants show no such enrichment **(Figure 3C; Supplemental Figure S23)**. Furthermore, no binding was observed in the mock experiment (immunoprecipitated with IgG) **(Figure 3C; Supplemental Figure S23)**. To further confirm that HY5 regulates genes associated with SGA and flavonoid biosynthesis, we developed promoter-reporter constructs using the promoters of *SlGAME9, SlGAME4*, and *SlMYB12*. These constructs were transiently expressed in tomato leaves of WT, SlHY5OX, and *SlHY5^CR^* plants. The histochemical GUS assay revealed a significant induction in GUS staining and expression in the overexpression lines of SlHY5 compared to WT plants. Conversely, no such change in GUS staining and expression was observed in *SlHY5^CR^* plants **(Supplemental Figure S24A-B)**. These results concluded that SlHY5 interacts with the SGA and flavonoid biosynthesis genes by binding to their promoters to regulate SGA and flavonoid biosynthesis, respectively.

### Light-dependent regulation of SGA and flavonoid biosynthesis

To study the influence of light on the expression of genes involved in SGA and flavonoid biosynthesis, we grew SlHY5OX and *SlHY5^CR^*seedlings in half-strength media in the continuous dark for 10 days, followed by exposure to different time intervals. Upon increasing the timing of light illumination, the expression of SGA biosynthesis genes (*GAME4/9/11/12/17* and *SlC5SD)* was significantly increased in WT and SlHY5OX seedlings. However, no such increase in expression was observed in *SlHY5^CR^* seedlings **(Figure 4A-F).** Additionally, the expression of flavonoid pathway genes (*SlCHS* and *SlFLS*) was evaluated and results indicated that light exposure enhanced transcript levels in SlHY5OX but not in *SlHY5^CR^* seedlings **(Supplemental Figure S25A-B).** These results suggested that light-dependent expression of SGA and flavonoid biosynthesis genes is dependent on HY5 in tomato.

**Figure 4.**
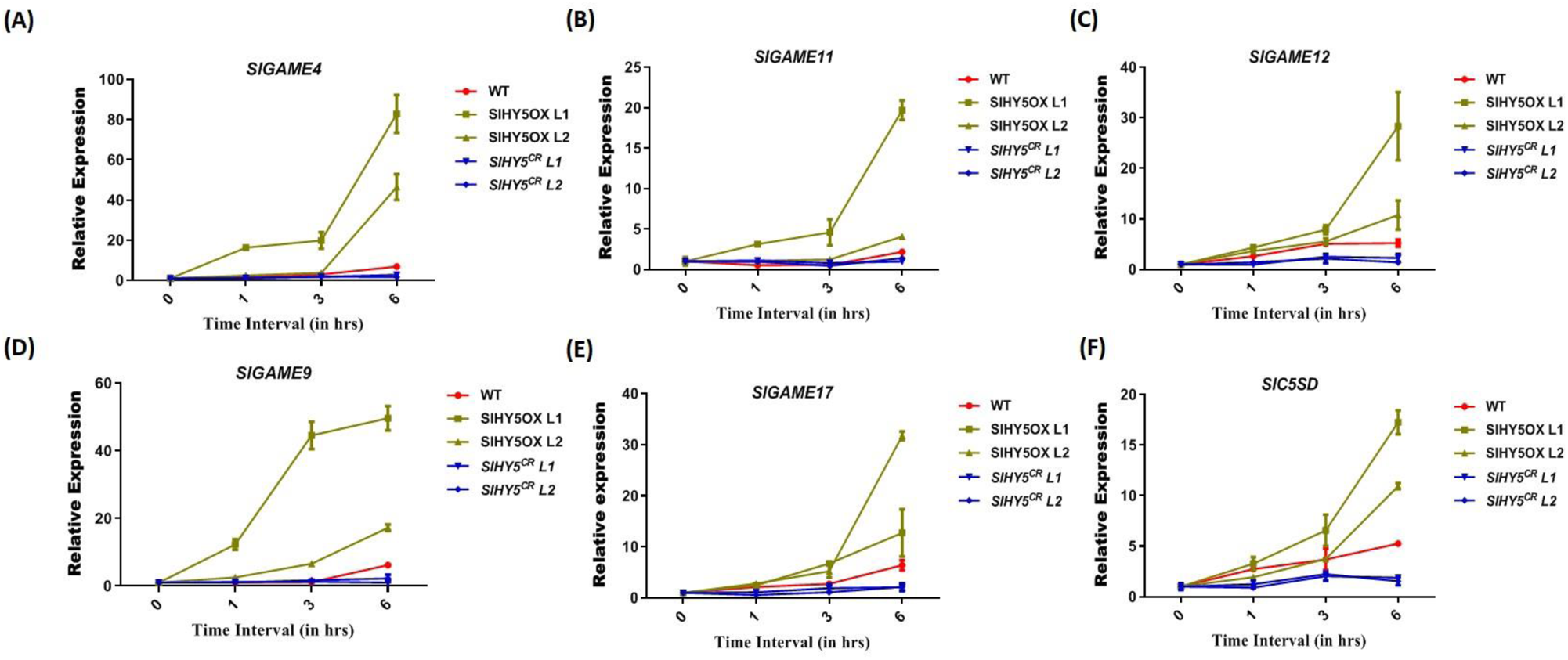
Light-mediated regulation of SGA in overexpression and CRISPR/Cas9-mediated knockout of *SlHY5*. **A-F,** Time dependent qRT-PCR expression analysis of SGA pathway genes (*SlGAME4, SlGAME11, SlGAME12, SlGAME17, SlC5SD* and *SlGAME9*) in 10-days-dark grown WT, SlHY5OX and *SlHY5^CR^*seedlings of tomato and the seedlings are subsequently transferred to light for 1, 3, and 6 hours. The statistical analysis was performed using two-tailed Student’s t-tests. The data are plotted as means ± SD (n=3). The error bars represent standard deviations. The asterisks indicate significant difference, *P< 0.1; **P< 0.01; ***P< 0.001.

All above-described results suggested the regulation of SGA and flavonoid biosynthesis genes by SlHY5 in tomato. For further validation, the accumulation of SGAs (α-tomatine and dehydrotomatine) and flavonoids were measured in the leaves and green fruits of WT, SlHY5OX and *SlHY5^CR^*plants **(Figure 5A).** SGAs (α-tomatine and dehydrotomatine) content decreased by 29.8% and 17.9% in *SlHY5^CR^* in leaves, while 66.2% and 42.8% in fruits compared to WT **(Figure 5B).** A significant increase in SGA (α-tomatine and dehydrotomatine) content was observed in the SlHY5OX plant compared to the leaf and fruit tissues of WT and *SlHY5^CR^*. The accumulation of major flavonoids (kaempferol, quercetin, rutin, and CGA) in SlHY5OX lines was higher, while it was lower in *SlHY5^CR^* lines in 10-day-old seedlings and fruit tissue **(Figure 5C; Supplemental Figure S26A-B)**.

**Figure 5.**
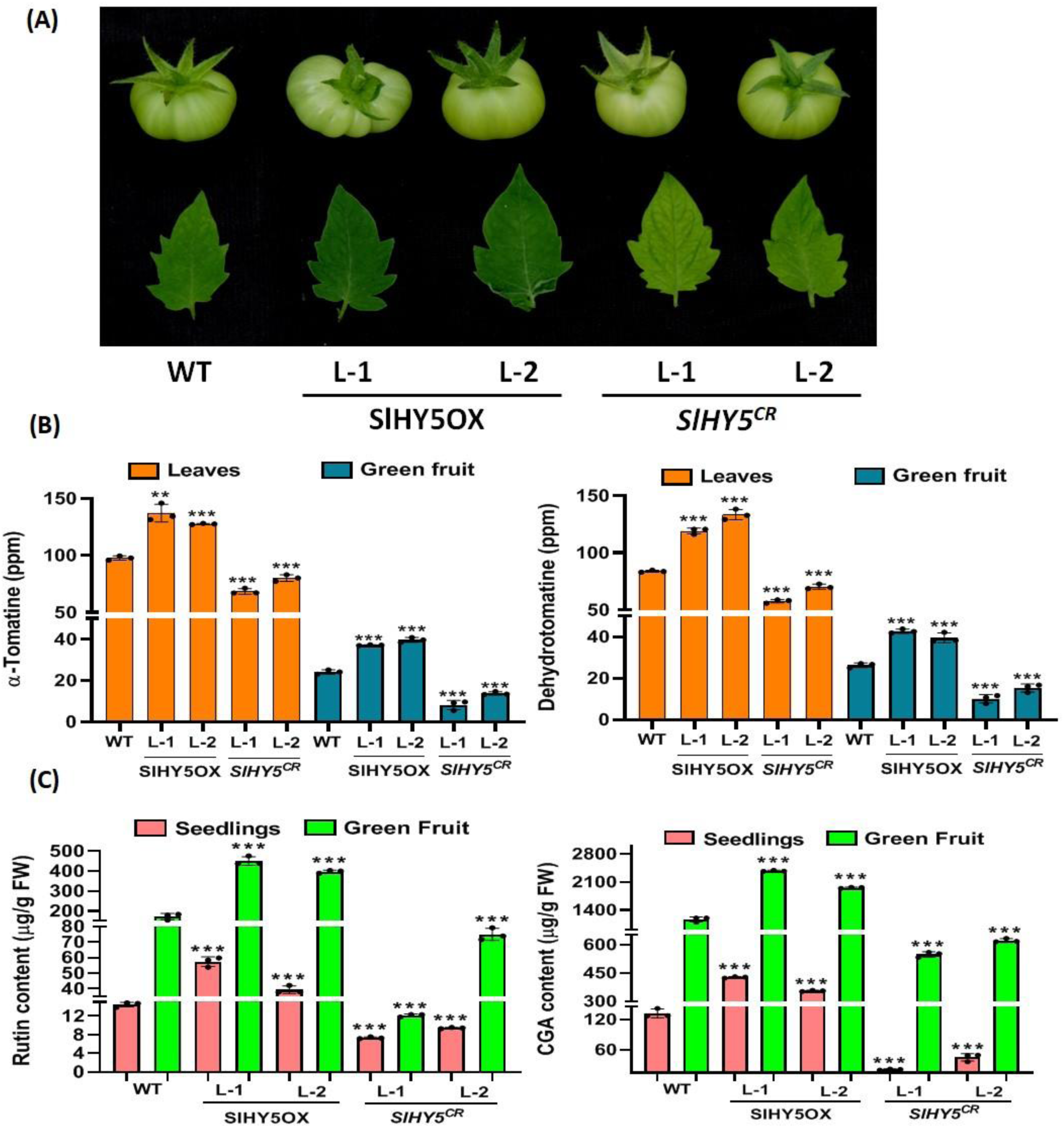
SlHY5 modulates SGA and flavonoid levels in tomato transgenic lines. **A,** Representative photograph of 1-months-old mature tomato leaf and green fruit being used for study. **B,** Quantitative estimation of SGAs (α-tomatine and dehydrotomatine) in 1-months-old leaf and green fruit of SlHY5OX and *SlHY5^CR^* tomato plants through using LC-MS. **C,** Quantification of flavonol (Rutin and CGA) content in 10-days-old seedlings and green fruit of SlHY5OX and *SlHY5^CR^* lines of tomato plants using HPLC. The statistical analysis was performed using two-tailed Student’s t-tests. The data are plotted as means ± SD (n=3). The error bars represent standard deviations. The asterisks indicate significant difference, *P< 0.1; **P< 0.01; ***P< 0.001.

### HY5-dependent differential response towards fungal infection

Previous reports suggested that SGA and flavonoid accumulation is related to biotic stress tolerance like fungal infection (Al Aboody et al., 2020; Bailly, 2021). Therefore, we analysed the response of SlHY5OX and *SlHY5^CR^*plants with the modulated accumulation of SGA and flavonoid content towards *Alternaria solani*. The leaves of 60-day-old plants of WT, SlHY5OX, and *SlHY5^CR^* were infected with the fungus. Analysis suggested that infection rate was higher in *SlHY5^CR^*lines while SlHY5OX lines have minimum infection compared to the WT **(Figure 6A and Supplemental Figure S27)**. This was validated through CFU (colony forming unit) count after infection, which showed a higher number of viable cells in *SlHY5^CR^* lines and a lower number of viable cells in SlHY5OX plants compared to WT. Here, CFU is directly correlated with the number of viable cells, showing the severity of the infection. Therefore, infection was higher in *SlHY5^CR^* lines and showed a sensitive phenotype, and SlHY5OX lines showed a lower infection rate and tolerant phenotype compared to WT **(Figure 6B).** As stress conditions induce ROS accumulation in plants, we estimated ROS in leaves of SlHY5OX and *SlHY5^CR^* lines after fungal treatment. DAB staining showed higher ROS levels (as brown precipitate) in *SlHY5^CR^* and a lower in SlHY5OX plants compared to the WT under biotic stress conditions **(Figure 6C).** Furthermore, NBT staining results suggested a lower accumulation of O^2-^ radical in SlHY5OX lines, while a higher accumulation was found in *SlHY5^CR^* lines compared to WT **(Supplemental Figure S28).** Similarly, fungal bioassays were conducted on both green and red fruits **(Figure 7A)**. The analysis revealed that fruits from *SlHY5^CR^*plants were more susceptible to infection, whereas those from SlHY5OX plants exhibited resistance compared to the WT. To validate these findings, CFU counts were performed, demonstrating a lower number of viable cells in the fruits of SlHY5OX plants compared to *SlHY5^CR^*plants **(Figure 7B)**. According to prior research, oxidative stress induced by biotic stress leads to tissue damage and lipid peroxidation through the generation of free radicals (Sahu et al., 2022). To quantify lipid peroxidation and cell wall degradation in the fruits of SlHY5OX and *SlHY5^CR^* following fungal treatment, we measured malondialdehyde (MDA) content and glucose liberation, respectively. Under stress conditions, SlHY5OX lines exhibited lower MDA content and glucose liberation, whereas mutated plants showed higher accumulation of MDA and glucose liberation compared to the WT in both green and red fruits **(Figure 7C; Supplemental Figure S29A-B)**.

**Figure 6.**
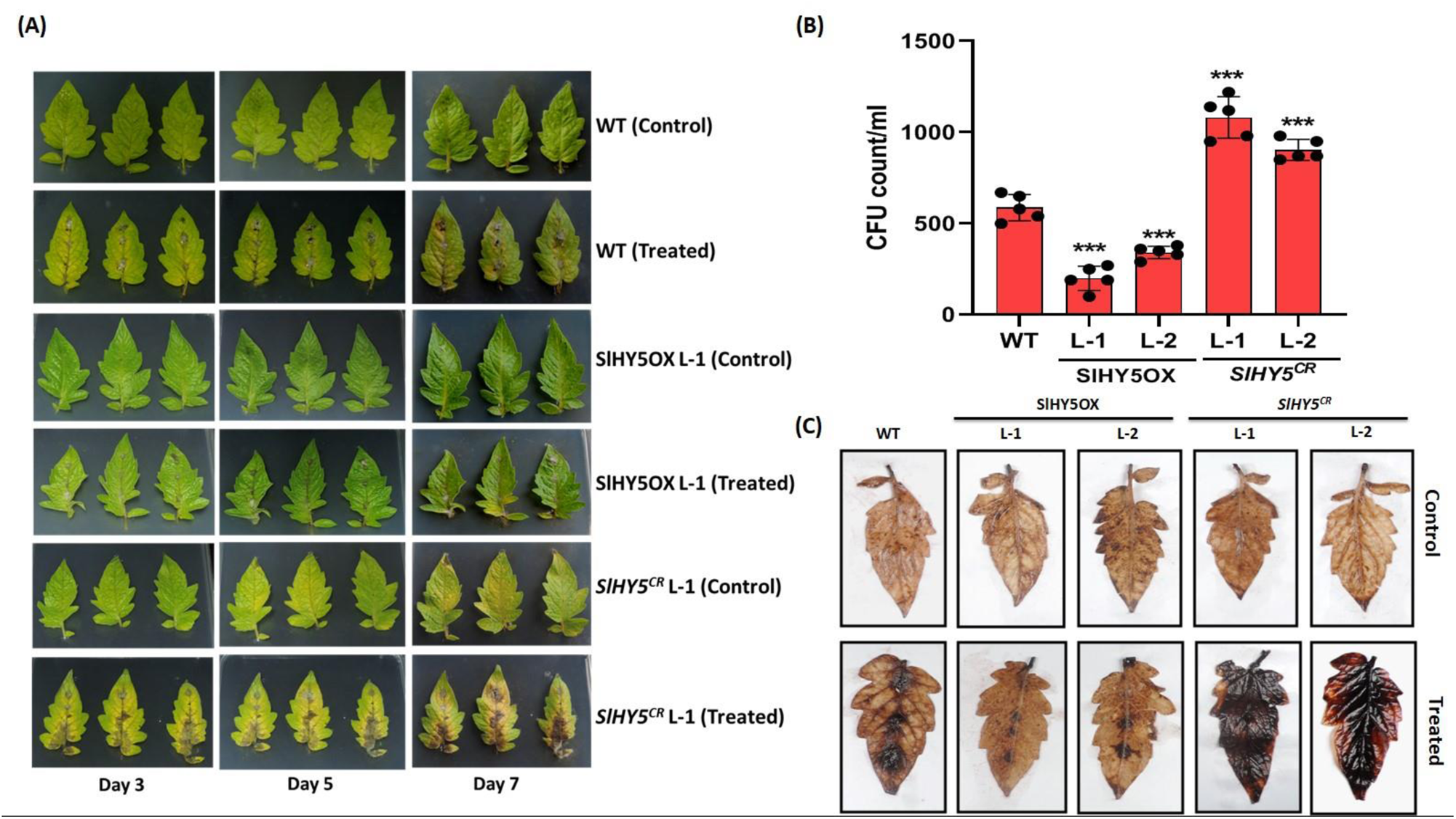
Fungal bioassay of tomato transgenic lines. **A,** Representative images of necrotic areas induced by *Alternaria solani* in detached tomato leaflets treated with 60 µl of conidial suspension. Leaflets spotted with 10mM MgSO_4_ with 0.4% gelatine only were included as the negative controls. **B,** CFU count of WT, SlHY5OX (L-1 and L-2) and *SlHY5^CR^* (L-1 and L-2) after infection. **C,** Staining of WT, SlHY5OX (L-1 and L-2) and *SlHY5^CR^* (L-1 and L-2) leaves with 3-3′diaminobenzidine (DAB) after fungal treatment. The statistical analysis was performed using two-tailed Student’s t-tests. The data are plotted as means ± SD (n=5). The error bars represent standard deviations. The asterisks indicate significant difference, *P< 0.1; **P< 0.01; ***P< 0.001.

**Figure 7.**
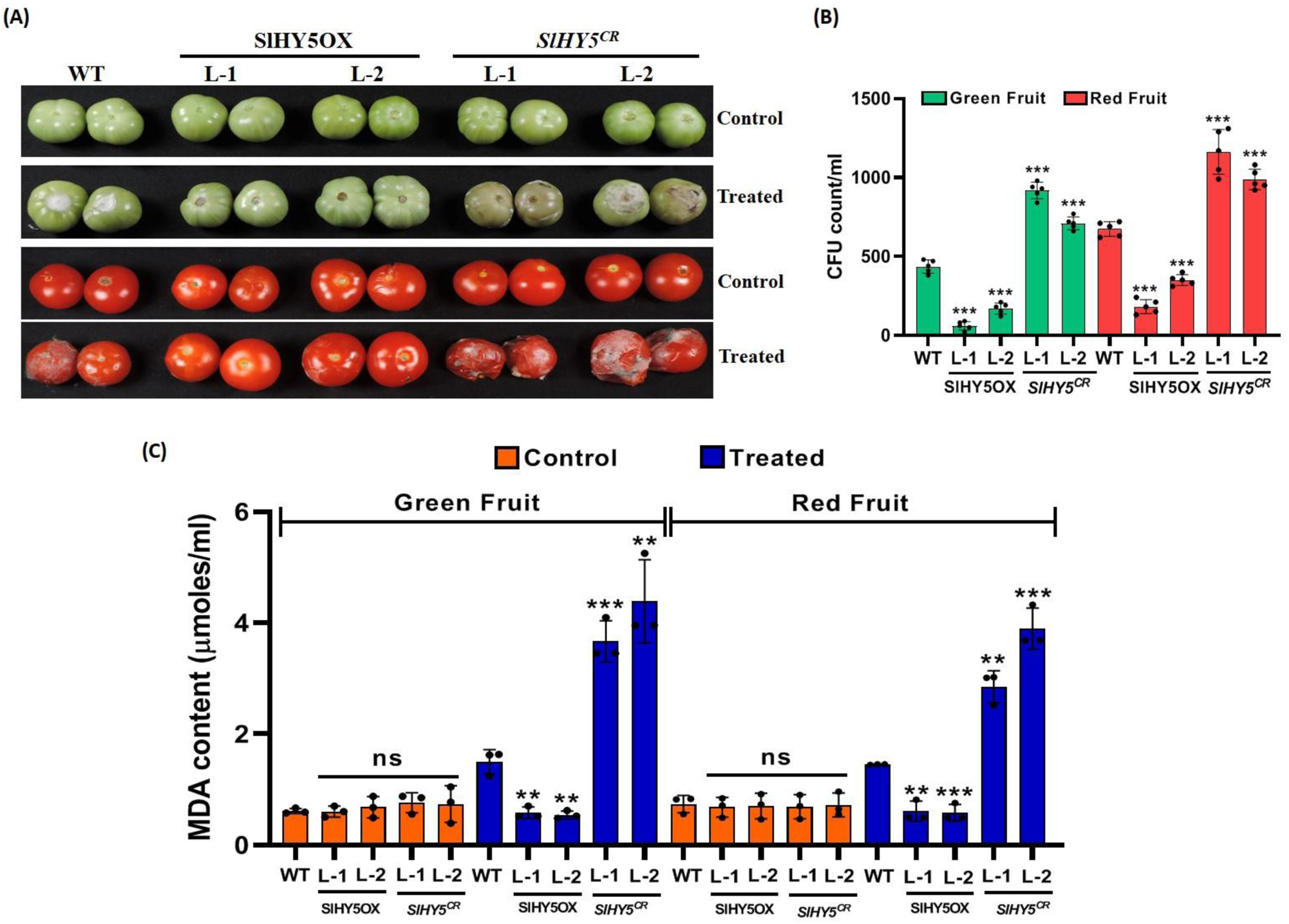
Fungal bioassay of tomato fruits. **A,** Representative images of infection areas induced by *Alternaria solani* in detached tomato fruits treated with 1ml of conidial suspension. The fruits spotted with 10mM MgSO_4_ with 0.4% gelatine only were included as the negative controls. **B,** CFU count of WT, SlHY5OX (L-1 and L-2) and *SlHY5^CR^* (L-1 and L-2) after infection in green and red fruits (n=5). **C,** Estimation of MDA content in WT, SlHY5OX (L-1 and L-2) and *SlHY5^CR^*(L-1 and L-2) fruits after fungal treatment. The statistical analysis was performed using two-tailed Student’s t-tests. The data are plotted as means ± SD (n=3). The error bars represent standard deviations. The asterisks indicate significant difference, *P< 0.1; **P< 0.01; ***P< 0.001.

Furthermore, fungal bioassays conducted on whole plants revealed a similar phenotype in leaves and fruits, with SlHY5OX lines displaying greater health than *SlHY5^CR^*plants and the WT **(Supplemental Figure S30)**. To investigate the role of SlHY5 in mediating light-dependent SGA and flavonoid biosynthesis as a mechanism of fungal tolerance, we conducted defense experiments on plants exposed to continuous light and darkness for a short period (48 hours). Our analysis indicated that SlHY5OX lines accumulated more SGA and flavonoid content in light conditions compared to darkness, contributing to tolerance against fungal attack **(Supplemental Figure S31A-C)**. To validate this experiment, we measured MDA content and found a lower amount of MDA in SlHY5OX lines in light-grown seedlings compared to those grown in darkness **(Supplemental Figure S31D)**. These results suggest that downregulation of SlHY5 leads to a higher accumulation of ROS, tissue damage and sensitive phenotype towards stress, whereas a higher accumulation of SlHY5 causes a lower accumulation of ROS, less tissue damage and tolerant phenotypic response in tomato under biotic stress due to modulated levels of SGAs and flavonoids.

## DISCUSSION

SGAs are known for their antinutritional properties; however, they play an important role in providing resistance to diseases and anti-herbivory responses in plants. Additionally, SGAs have been studied for their antifungal, antidiabetic, antiparasitic, antimicrobial, antibiotic, antiviral, and generally for anticancer properties (Heftmann 1983; Li et al., 2020; Lee et al., 2007; Milner et al., 2011; Caputi et al., 2018). Approximately, 100 SGAs are found in tomato, including α-tomatine and dehydrotomatine, which are accumulated in various plant organs like leaves, flower buds, and green fruit tissues (Itkin et al., 2011; Moco et al., 2006; Moco et al.,2007; Iijima et al., 2013; Li et al., 2020). Previous studies suggested that a group of JREs (jasmonate-responsive transcription factors of the ETHYLENE RESPONSE FACTOR) family, such as SlGAME9, is involved in the regulation of SGA regulation (Thagun et al., 2016; Cárdenas et al., 2016). The presence of LREs (light-responsive elements) in the SGA biosynthesis genes suggested the involvement of light in the SGA biosynthesis pathway. The silencing of light-associated transcription factors, SlHY5 and SlPIF3, using VIGS showed significantly altered SGA levels in tomato leaves compared to WT plants (Wang et al., 2018).

In this study, we have compared light-grown to dark-grown WT seedlings for the accumulation of SGA and flavonols. Analysis suggested a higher transcript level and accumulation of SGA and flavonols in light-grown seedlings, indicating the role of light signaling components in their biosynthesis **(Figure 1B-E and Supplemental Figure S1A-B)**. An increase in the expression of SGAs biosynthesis genes in dark-grown WT seedlings exposed to light at different time intervals led us to hypothesize that the SlHY5 may participate in SGA and flavonoid biosynthesis in tomato **(Figure 1G and Supplemental Figure S1C)**. In the presence of light, HY5 affects their putative targets by binding to the light-responsive elements (LREs) majorly with G-box, C-box, and T/G-box (ACGT-containing cis-element) (Lee et al., 2007; Song et al., 2008). It has been shown that the SGAs/flavonol/anthocyanin metabolic pathway genes also contain G-box in their promoters, suggesting the role of HY5 in plant secondary metabolism (Pan et al., 2009; Wang et al., 2018). Wang et al. (2018) used EMSA to validate the binding of SlHY5 with the promoters of GAME4, GAME1, and GAME17. The group used VIGS of *SlHY5* to functionally characterize its role in SGA metabolism and found that VIGS of HY5 led to a reduction of transcript abundance of SGA biosynthesis genes, leading to reduction in α-tomatine levels. Our *in-silico*, EMSA, ChIP-qPCR, and transient assay studies suggest an interaction of SlHY5 with promoters of additional genes associated with the SGA biosynthetic pathway due to the presence of G-box in the promoter of *SlGAME9/4/11/12/17/SlC5SD* and *SlMYB12* **(Figure 3 and Supplemental Figure S20-24)**. Furthermore, while Wang et al. (2018) used VIGS to functionally characterize HY5 in tomato, we developed SlHY5OX and *SlHY5^CR^* to validate the results and showed that, in addition to SGA biosynthesis, SlHY5 regulates biotic stress response in tomato.

Various plant species belonging to different families contain orthologs of HY5, responsible for photomorphogenesis. Some of these orthologs of HY5 have been shown to complement the function when overexpressed in *Arabidopsis hy5* mutant (Song et al., 2008). Rice ortholog OsZIP48 was also capable of recovering the lost phenotype of *hy5* mutant plants of *Arabidopsis* after complementation assay (Song et al., 2008; Burman et al., 2018). Subsequently, to discover whether SlHY5 acts as a functional ortholog of AtHY5 and NtHY5, we performed the complementation study. The results demonstrated that SlHY5 could restore the loss of phenotype (hypocotyls length, root length, rosette diameter, seed size, plant height) as well as transcript level. Accumulation of metabolites like chlorophyll, and flavonoids were also rescued in SlHY5:*hy5* compared to *hy5* mutant of *Arabidopsis* and tobacco **(Supplemental Figure S6-15)**. These findings validated that the complementation of *Arabidopsis* and tobacco *hy5* by SlHY5 regulates flavonoid biosynthesis as they share conserved functional homology with other plant species **(Supplemental Figure S4)**.

Further investigations were done to unveil the influence of SlHY5 in SGAs and flavonoid biosynthesis by generating the overexpression and CRISPR/Cas9-mediated knockout lines in tomato **(Supplemental Figure S16 and S17A)**. α-tomatine and dehydrotomatine are the best-known alkaloids present in tomato, majorly accumulated in the green stages of plants like green fruits, leaves, etc., and their level decreases at the red stage of development (Moco et al., 2007). Accordingly, we have carried out the analysis of SGAs content and its pathway gene expression in leaves and green fruits of overexpression and mutant plants. The elevated transcript level in overexpression and lower level in *SlHY5^CR^*plants compared to WT was observed in leaves and fruits **(Figure 2C)**. The SGAs accumulation was also affected in SlHY5OX and *SlHY5^CR^* lines as the SGA content was significantly higher and lower in these plants, respectively **(Figure 5B).** Time-dependent regulation of SGAs pathway genes in SlHY5OX and *SlHY5^CR^* lines, which are dark-grown seedlings, towards light exposure indicated an increase in expression of genes in SlHY5OX lines compared to WT, but no changes were found in *SlHY5^CR^*plants, suggesting the involvement of light in SGAs biosynthesis **(Figure 4).** Sequentially, we carried out the experiment in 10-day-old seedlings to analyze the expression of phenylpropanoid pathway-related genes in SlHY5OX and *SlHY5^CR^* plants. Analysis suggested the higher expression of overexpression compared to WT, while the opposite results were found in mutated lines. A similar result was also observed in fruit tissue **(Supplemental figure S18B-D)**. Time-dependent studies also suggested the involvement of SlHY5 in flavonoid biosynthesis as maximum induction was found in 3 h in overexpression, but no significant change in *SlHY5^CR^*lines **(Supplemental Figure S25)**.

During plant growth and development, various transcription factors, including HY5, and their specific targets play an important role (Gangappa et al., 2016; Yang et al., 2018; Zhang et al., 2020). Inhibition of auxin signaling affects the hypocotyl elongation by HY5; also, plant height was reduced due to the reduction in the cell size in OsbZIP48 overexpression lines which is an ortholog of AtHY5. Additionally, in many aspects of plant physiology, flavonoids play an important role to regulate root and shoot development by influencing polar auxin transportation in various conditions (Peer and Murphy 2007; Burman et al., 2018; Qian et al., 2023). We perform morphometric analysis of developed SlHY5OX and *SlHY5^CR^*lines since our data revealed the involvement of SlHY5 in regulating SGA/flavonoid biosynthesis. We measured the phenotype of 10-day-old light-grown plants of SlHY5OX and *SlHY5^CR^* plants and the results suggested that variations in phenotype were due to alteration in SGAs (α-tomatine and dehydrotomatine) and flavonoid content in plants. Briefly, the length of hypocotyls was shorter in overexpressed lines, but it was longer in mutant lines compared to WT **(Figure 2A).** One-month-old plants showed a significant difference in their height **(Supplemental Figure S17B)**. This data concluded the involvement of SlHY5 in cell elongation leads to changes in phenotype.

According to previous studies, various biotic factors (viruses, bacteria, fungi) are responsible for yield loss in tomato and among them fungi cause the most destructive foliar diseases in most of the Solanaceae plants (Jones et al., 2014; Jones and Perez, 2019; Neergaard et al., 1945; Chaerani et al., 2006; Attia et al., 2020). Furthermore, mutation in HY5 enhances the ER stress tolerance and increases the expression of unfolded protein response (UPR), showing the negative regulation of UPR by light (Nawkar et al., 2017). HY5 expression is regulated by SPA kinases to regulate root thermomorphogenesis, and other than HY5, many transcription factors (PIFs, BZR1, MYC) are involved in plant growth and defence by transcriptional feedback (Lee et al., 2021; Koene et al., 2022). Also Panda et al. (2022) suggested the mechanistic basis of hormone signaling in regulating SGA to balance the growth and defence. In their study, Panda et al. (2022) investigated the hormonal regulation of SGAs metabolism in the context of growth-defense trade-off. Through genetic and biochemical analyses of JA and GA pathways, along with in vitro experiments, they explored the interplay between these hormone pathways in regulating SGAs metabolism. Their findings revealed that reduced JA levels decrease SGA production, while low GA levels lead to increased SGA accumulation. The study concluded that MYC1 and MYC2 transcription factors mediate the crosstalk between JA and GA by activating SGA biosynthesis and GA catabolism genes. Further support for the JA/GA crosstalk during SGA metabolism was provided by chemical and fungal pathogen treatments.

As, the accumulation of SGAs and flavonoids provides defense from biotic stress (such as fungi), we performed the pathogenicity test in the leaves, fruits and whole plants of tomato, suggesting the *SlHY5^CR^* lines are more susceptible to disease compared to the WT which might be due to lesser accumulation of SGA and flavonoid **(Figure 6A-7A and Supplemental figure S27 and S30)**. Biotic or abiotic stresses increase the accumulation of Reactive Oxygen Species (ROS) in plants (Jalmi and Sinha, 2015). These ROS radicals play a role in signal transduction in plants, but the excess level results in membrane disruption and nucleic acid damage eventually, causing lipid peroxidation and tissue damage (Mittler, 2017; Choudhury et al., 2017; Singh et al., 2019; Sahu et al., 2022). Increased DAB/ NBT staining as well as MDA content and glucose liberation were observed in *SlHY5^CR^* lines, which suggests a higher ROS production in these plants compared to WT **(Figure 6C-7C and Supplemental Figure S28-S29)**. This might be a possible explanation for the sensitivity of *SlHY5^CR^* under biotic stress conditions. Moreover, SlHY5OX plants have a lesser accumulation of ROS than WT, which showed clear evidence that SlHY5 positively regulates tolerance toward fungal infection **(Figure 6C and Supplemental Figure S28)**. Light-dependent regulation of SGAs and flavonoids as a fungal tolerance mechanism also indicated fungal infection was higher in dark condition compared to light due to lower accumulation of SGA and flavonoids in dark compared to light **(Supplemental Figure S31)**.

Based on our results and previous studies discussed earlier, we propose a model illustrating light-dependent SGA and flavonoid biosynthesis mediated through SlHY5 **(Figure 8)**. The presence of G-BOXES on the promoter region of genes indicating the regulation of SGA/flavonoid biosynthesis by light. The regulation by SlHY5 leads to modulating plant growth and development such as plant height, hypocotyl length, etc. Furthermore, increased SGA and flavonoid content suggest the positive regulation of SlHY5 in their biosynthesis and defense mechanism against fungal infection, respectively. Thus, this study provides an imminent molecular strategy to develop transgenic tomato, having significantly modulated SGAs and flavonoids through manipulating the SlHY5 gene of tomato.

**Figure 8.**
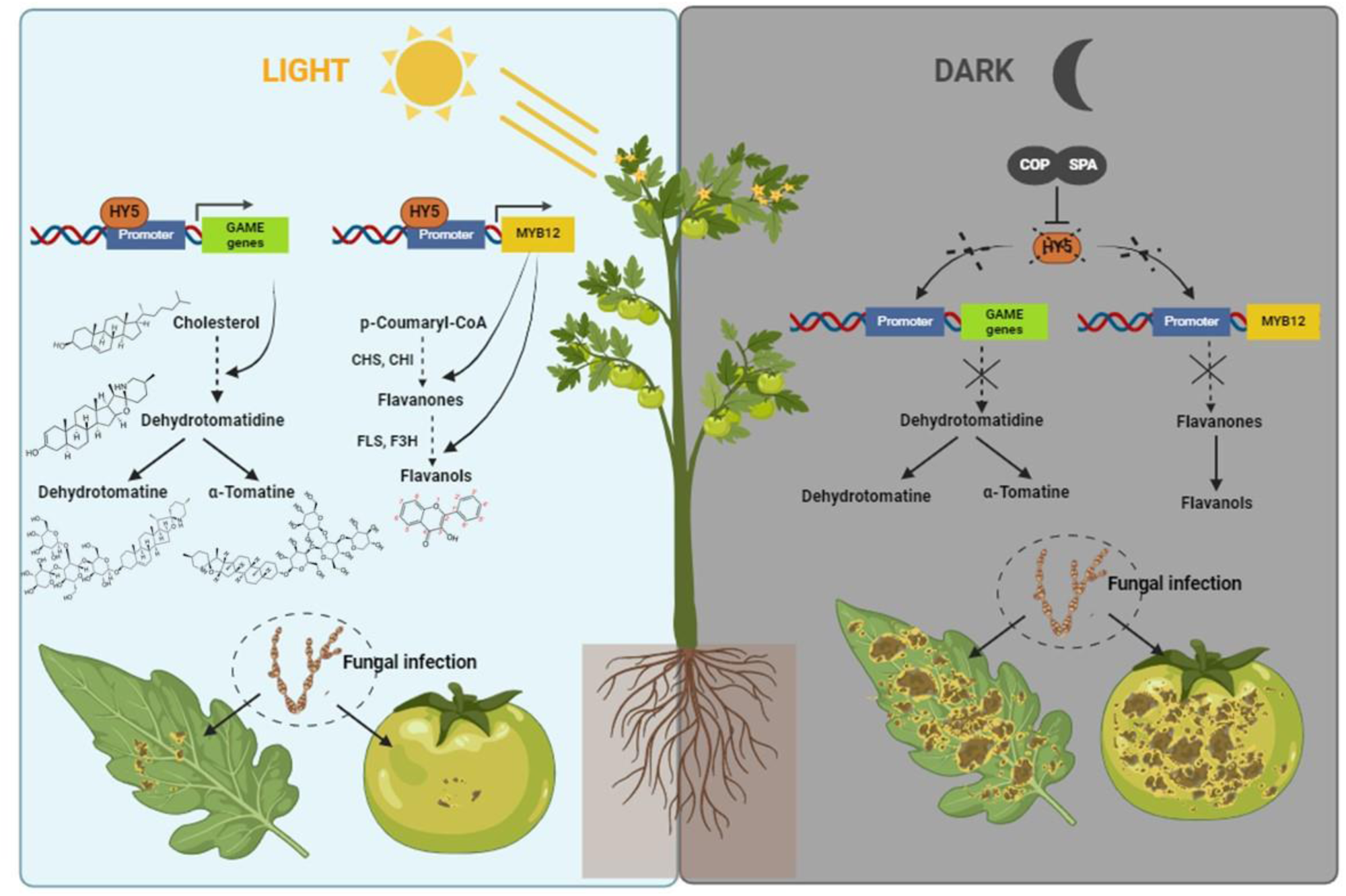
Proposed working model showing the role of SlHY5 in SGA and flavonol accumulation in tomato. We propose that presence of light activates SlHY5 which upregulate the expression of SGA and flavonoid biosynthesis genes, leading to enhanced accumulation of SGA as well as flavonoid biosynthesis. SlHY5 regulates SGA and flavonoid biosynthesis via binding to the G-boxes in the promoters of SGA and flavonoid regulatory as well as structural genes. Higher accumulation of secondary metabolites and higher antioxidant activity thus increase biotic stress tolerance ability. On the contrary, absence of light exhibits the opposite effect. Arrowheads and tees indicate positive and negative regulation, respectively.

## MATERIALS AND METHODS

### Plant material and growth conditions

Cotyledonary leaves of tomato (cv. Pusa Ruby) seedlings were used for raising the transgenic plants. Tomato plants were grown in the glass house at 24°C±2°C and 16 h/8 h light-dark photoperiods for harvesting of tissues belonging to different developmental stages. For germination, seeds were surface sterilized using 70% EtOH for 2-4 min and then dipped in 50% bleach for 15-20 min and washed repeatedly with autoclaved Milli-Q water and placed on half-strength MS medium (Hi-Media) containing 1.5% sucrose, pH 5.72–5.8. After stratification for 3 days in the dark, the plates were transferred to a culture room with 16 h/8 h light-dark photoperiods, 24°C temperature. For dark treatment, AtMYB12 overexpressing seedlings (developed earlier in our lab by Pandey et al., 2015) were grown in light for 10 days and then harvested after 24 h and 48 h of dark exposure for gene expression as well as phytochemical analysis. Samples were frozen in the liquid N_2_ and kept in a −80°C deep freezer until use. The experiment was performed by using three independent replicates. *Arabidopsis* (Col-0) and *Nicotiana tabacum cv. Petit Havana* was used as the wild-type plant in the complementation study. Seeds were surface sterilized and placed on a one-half-strength MS medium containing 1.5% Sucrose. After stratification for 2 days at 4°C in the dark, seeds were transferred to a growth chamber under controlled conditions of 16-h-light/8-h-dark photoperiod cycle, 22°C temperature, 150 to 180 μmol m^−2^s^−1^ light intensity, and 60% relative humidity for 10 days unless mentioned otherwise.

### Identification and sequence analysis of SlHY5

Identification of the ortholog of AtHY5 in tomato was performed at National Center for Biotechnology Information (http://www.ncbi.nlm.nih.gov/BLAST) using BLASTn. The sequence of SlHY5 was retrieved, which showed 90% similarities with AtHY5 and NtHY5. Analysis of amino acid sequence for the leucine-rich bZIP Zinc finger domain and VPE domain was done using a Multalin tool. To ensure that identified SlHY5 protein belonged to the bZIP family, a conserved domain database (CCD) was used. Using the PlantCare tool (Lescot et al., 2002), the promoter sequence of the *SlMYB12* gene (∼1.5 kb upstream of translation start site) was retrieved, which was further utilized for the screening of light-responsive cis-elements (LREs). National Center for Biotechnology Information BLASTn tool was used to identify the homologs of AtHY5 that belong to different families. The genes retrieved were analyzed for the presence of the requisite domains using InterProScan (http://www.ebi.ac.uk/Tools/pfa/iprscan/; Quevillon et al., 2005). The evolutionary history was inferred by using the maximum likelihood method and JTT matrix-based model. The tree with the highest log likelihood (-6071.97) is shown. Initial tree(s) for the heuristic search were obtained automatically by applying neighbor join and BioNJ algorithms to a matrix of pairwise distances estimated using the JTT model, and then selecting the topology with superior log likelihood value. This analysis involved 35 amino acid sequences. Evolutionary analysis was conducted in MEGA11.

### Plasmid construction and generation of *Arabidopsis*, tobacco, and tomato transgenic lines

For overexpression of SlHY5, the SlHY5 gene was amplified from the cDNA library prepared from 10-day-old tomato seedlings using gene-specific primers through PCR. The full-length ORF of the SlHY5 cDNA was cloned in the binary vector pBI121 (Clontech, USA), under the control of cauliflower mosaic virus 35S promoter. The construct was transferred into *Agrobacterium tumefaciens* strain GV3101 and used to transform *Arabidopsis* (Col-0) by the floral dip method (Clough and Bent, 1998) and tobacco and tomato plants by leaf disk method (Horsch et al., 1985). Several transgenic tomato lines were selected based on the expression analysis. Seeds were harvested, surface sterilized, and plated on a solid half-strength MS medium supplemented with 100 mg/l kanamycin. Antibiotic-resistant plants were transferred to the glass house and grown until maturity.

CRISPR-Cas9 technology was employed, for the development of mutant plants of SlHY5. A 20 bp gRNA was selected from the identified exonic region. gRNAs of SlHY5 were cloned into the binary vector pHSE401 using the BsaI restriction site (Xing et al., 2014). This vector contains Cas9 endonuclease-encoding gene under dual CaMV35S promoter as well as genes encoding neomycin phosphotransferase and hygromycin phosphotransferase as selection markers. All the constructs were sequenced from both the orientations using plasmid-specific forward and vector reverse primers and that transferred to GV3101 and then transformed to tomato. To perform the transient expression, genomic DNA of WT tomato seedling was isolated and PCR amplified the 1.5 kb promoter region (upstream from translation initiation codon ATG) using High Fidelity enzyme mix (Fermentas, USA) and gene-specific primers. After cloning into pBI121, the *SlGAME9pro*::GUS, *SlGAME4pro*::GUS, and *SlMYB12pro*::GUS were introduced into *Agrobacterium* (strain GV3101) and then infiltrated in leaves of WT, SlHY5OX and *SlHY5^CR^* plants.

### Expression analysis

For quantitative PCR, total RNA was isolated and treated with TURBO DNAase (Ambion), and 1 μg was used for reverse transcription employing the Revert AidH plus First Strand cDNA Synthesis Kit (Fermentas) as per the manufacturer’s instructions. The cDNA was 20 times diluted with nuclease-free water, and for quantitative PCR, 2 μl of cDNA was used as a template using Fast SYBR Green Mix (Applied Biosystems) in a Fast 7500 Thermal Cycler instrument (Applied Biosystems). The expression was normalized using internal control as Actin and analyzed through the comparative ΔΔ^CT^ method.

### Total flavonol quantification

Ten-day-old seedlings (for tomato as well as *Arabidopsis*) were used for flavonol estimation. Extraction was done in 1 ml of 80% methanol at 4 °C for 2 h shaking. The mixture was centrifuged at 12,000 g for 12 min. The supernatants (0.5 ml) were taken following the addition of 2 ml methanol and subsequently mixed with 0.1 ml of aluminium chloride (10% water solution), 0.1 ml of potassium acetate (1 M), and 2.8 ml of MQ water. After incubation for 30 min, the absorbance was recorded at 415 nm. The calibration curve was prepared using quercetin as the standard. The total flavonol content was calculated as the equivalent of quercetin used as the standard.

### Extraction and quantification of flavonols

Estimation of flavonols was done either as aglycones or as flavonol glycosides by preparing acid-hydrolyzed or non-hydrolyzed extracts respectively. For extraction, the plant material (300 mg) was ground into the fine powder in liquid N_2_ and transferred the sample in 80% methanol overnight at room temperature with brief agitation. An equal amount of 6N HCl was added to the extract for hydrolysis at 70°C for 40 min subsequently an equal amount of methanol was added to prevent the precipitation of the aglycones (Jiang et al. 2015). Filtration of extracts was done through a 0.2 mm filter (Millipore, USA) before HPLC. For non-hydrolyzed extracts, samples were extracted as described previously (Pandey et al., 2014). Breeze 2 software (Waters) was used for the quantification of various metabolites through HPLC. For the mobile phase, a gradient was prepared from 0.05% (v/v) ortho-phosphoric acid in HPLC-grade water (component A) and methanol (component B). Prior to use, these components were filtered through 0.45-mm nylon filters and deaerated in an ultrasonic bath. The gradient from 25 to 50% B for 0–3 min, 50 to 80% B for 3–18 min, 80 to 25% B for 25 min, and 25% B for 30 min was used for conditioning of the column with a flow rate of 1ml/min. All the samples were analysed by HPLC–PDA with a Waters 1525 Binary HPLC Pump system comprising PDA detector following the method developed by Niranjan et al., 2011.

### Chlorophyll estimation

Chlorophyll content was estimated by taking the 200mg tissue of 10-day-old WT and transgenic seedlings. Sample was crushed in liquid N_2_ and transferred in 2 ml (80 %, v/v) of chilled acetone (Arnon 1949). Homogenized tissues were centrifuged for 10 min at 8000g. Supernatant was collected and absorbance was recorded at 663,652, 645, 510 and 480 nm. Total chlorophyll content was calculated in mg g^−1^ fresh wt using formula of Maclachlan and Zalik (1963). Total chlorophyll= [34.5(A652)*V]/d *1000*W.

Where, V=volume of the final extract (ml), W= weight of the plant sample (g) and d=width of the cuvette(1cm).

### Antioxidant activity

To determine the antioxidant activity, DPPH (1, 1-diphenyl-2 picrylhydrazyl free radical) assay was used. In brief, DPPH solution (0.1 mM) was prepared in methanol and the initial absorbance was measured at 515 nm which was used throughout the assay. Samples were aliquoted (50–100 μl) and make up with DPPH solution to final volume of 1.5ml as described in Wong et al., (2006). The samples were incubated at room temperature for 30 min. The absorbance was recorded at 515 nm using Ultraspec 3000 UV/Vis spectrophotometer (Pharmacia Biotech Ltd., Cambridge, CB4, 4FJ, UK). Trolox was used to make the calibration curve using different dilutions. The antioxidant capacity based on the DPPH free radical scavenging ability of the extract was expressed as mM Trolox equivalents per gm of plant material on fresh weight basis.

### Expression and purification of recombinant SlHY5

pET-28b(+) expression vector (Novagen, Germany) was used for cloning of complete ORF of SlHY5. The construct was transformed into *E. coli* BL21 (DE3) pLysS (Invitrogen, USA) for prokaryotic expression which was induced by IPTG. Ni-NTA column (Nucleopore, India) was used to purify recombinant 6X-His-tagged HY5 and the eluate was quantified through Bradford assay.

### Electrophoretic mobility shift assay

Labelling of probes with digoxigenin was done using the 2nd generation DIG Gel Shift EMSA kit (Roche, USA) as per manufacturer’s instructions. Labelled probes were incubated for 30 min at 21°C in binding buffer [100 mM HEPES (pH 7.6), 5 mM EDTA, 50 mM (NH_4_)_2_SO_4_, 5 mM DTT, Tween 20, 1% (w/v), 150 mM KCl] with or without recombinant protein. To check specific binding, unlabelled probes were added to the reaction mixture in increasing concentrations. The binding reaction was resolved on a 6% polyacrylamide gel in 0.5X TBE buffer (pH 8.0) and was semi-dry blotted (Transblot, BIO-RAD, USA) onto a positively charged nylon membrane (BrightStar, Invitrogen, USA) followed by UV cross-linking. The membrane was finally incubated with CSPD chemiluminescent solution and exposed to X-ray blue film (Retina, India).

### Chromatin immunoprecipitation

Ten-day-old seedlings (2.5 g) were cross-linked with 30 mL of 1% formaldehyde in a vacuum for 20 min. A total of 2.5 ml of 2 M Glycine was added to stop the cross-linking. After rinsing seedlings with water, tissues were ground with liq-N_2_ and resuspended with 20 mL of EB I [0.4 M sucrose, 10 mM Tris-HCl, pH 8, 10 mM MgCl_2_, 5 mM beta-mercaptoethanol, 0.1 mM phenylmethylsulfonyl fluoride (PMSF), and protease inhibitor; Sigma], then filtered through cell strainer (Corning). The filtrate was centrifuged at 4000 rpm at 4^0^ C for 30 min. The pellet was resuspended in 1 mL of EB II (0.25 M sucrose, 10 mM Tris HCl, pH 8, 10 mM MgCl_2_, 1% Triton X-100, 5 mM beta-mercaptoethanol, 0.1 mM PMSF, and protease inhibitor) and centrifuged at 14,000 rpm and 4^0^ C for 10 min. The pellet was resuspended in 300 mL of EB III (1.7 M sucrose, 10 mM Tris-HCl, pH 8, 0.15% Triton X-100, 2 mM MgCl_2_, 5 mM beta-mercaptoethanol, 0.1 mM PMSF, and protease inhibitor) and loaded on top of an equal amount of clean EB III, then centrifuged at 14,000 rpm for 1 h. The crude nuclear pellet was resuspended in nuclear lysis buffer (50 mM Tris-HCl, pH 8.0, 10 mM EDTA, 1% SDS, and Complete protease inhibitor; Roche) and sonicated with a Branson sonifier (VWR) to achieve an average fragment size of 0.3 to 0.8 kb. The sonicated chromatin was centrifuged, and the insoluble pellet was discarded. The soluble chromatin solution was diluted 10-fold with ChIP dilution buffer (1.1% Triton X-100, 1.2 mM EDTA, 16.7 mM Tris-HCl, pH 8.0, and 167 mM NaCl), then after preclearing with protein A–Sepharose beads (Sigma-Aldrich), HY5 antibody (AS12 1867; Agrisera) was added to 1 mL of chromatin solution and incubated overnight at 4^0^C. The immunocomplexes were extracted by incubating with 100 mL of 50% protein A–Sepharose beads for 1 h at 4^0^C. After several washes, immunocomplex was eluted twice from the beads with 250 mL of elution buffer (1% SDS and 0.1 M NaHCO_3_), then reverse cross-linked with a final concentration of 200 mM NaCl at 65^0^ C for 12 h (overnight). After removing all proteins by treating with proteinase K, DNA was purified by phenol-chloroform extraction and followed by ethanol precipitation. The pellet was resuspended with 50 mL of 0.13 TE (10 mM Tris-EDTA, pH 7.5) with RNase A (0.1 mg/mL) and used for probe synthesis or PCR analysis.

### Transient expression assay

All the constructs (GUS gene under the control of *SlGAME9, SlGAME4 and SlMYB12* promoters), were transformed into *Agrobacterium tumefaciens* (strain GV3101) and used for agro-infiltration in four-week-old tomato leaves. *A. tumefaciens* strains were resuspended in 5 ml of LB (Luria broth) after 24 h of incubation at 28 °C, 150 rpm. The secondary inoculations were performed in 50 ml of solution (49 ml of LB and 1 ml of 0.1 M MES), and the cultures were incubated at 28 °C at 150 rpm overnight in induction medium (10 mM MgCl_2_, 10 mM MES (pH 5.6), 150 μM acetosyringone) and incubated for 2 h at 28 °C. The cultures were then diluted to an absorbance at 600 nm of 0.5 and injected into tomato leaves using a blunt-end 1 ml syringe. The plants were placed under constant light (∼70 μmol photons per m^2^ per s) for 48–60 h before the GUS assays were performed in infiltrated leaves.

### Histochemical GUS staining

The GUS staining was performed using a previously described method (Jefferson., 1989) with slight modifications. Leaves of WT and transgenic lines were immersed in a solution containing 100 mM sodium phosphate buffer (pH 7.2), 10 mM EDTA, 0.1% Triton X-100, 2 mM potassium ferricyanide, 2 mM potassium ferrocyanide and 1 mg ml−1 5-bromo-4-chloro-3-indolyl-β-D-glucuronide at 37 °C for overnight. The chlorophyll was removed by incubation and multiple washes using 70% ethanol and the pictures were taken for the GUS staining using camera.

### Analysis of SGAs though LC-MS

Tomato leaves (300 mg) were frozen and ground in liquid nitrogen. The ground powder derived was treated with 900 μl of 80% methanol (v/v). The mixture was vortexed for 30 sec, followed by its sonication for 30 mins at room temperature. After sonication, it was again vortexed for 30 sec and centrifuged at 20,000xg for 10 mins. The supernatant obtained was filtered through a 0.22 μm aperture PVDF membrane filter. The above process for extraction was repeated thrice. The final filtrate thus extracted was lyophilized. The lyophilized samples were reconstituted in 1 ml solution of 80% methanol (v/v). The reconstituted mixture was vortexed again and centrifuged at 20000xg for 10 mins. The final supernatant was transferred to an autosampler glass vial and subjected to LC-MS analysis. α-tomatine and dehydrotomatine were identified in the samples by comparing with the retention times and mass spectra of the corresponding standard compounds, which were earlier detected and analyzed by the same instrument (Cárdenas et al., 2016).

### Pathogenicity Test

The excised leaves were surface disinfected by dipping in 70% ethanol and sterile MQ for 30 seconds each and placed onto blotting paper to dry. Leaves were then placed in 0.8% agar plates. For green and red fruits, detached fruits were surface sterilized using 70% ethanol and sterile MQ for few seconds and then kept the fruits in sterilized flasks covered with cotton plug. Leaves and fruits were inoculated with conidial suspension of *Alternaria solani* having 1×10^6^ conidia/ml suspension and incubated at room temperature for the appearance of symptoms. Suspension was prepared in 10 mM MgSO_4_ containing 0.4% gelatine. This suspension (without conidia) was used as control. Pictures were taken 3, 5, and 7 days after application of suspension (5 and 10 days in case of red and green fruits, respectively). To measure the CFU, fungal suspension culture was spread to RBA (Rose Bengal Agar) plates and then calculated by the formula

> CFU/ml = (Number of colonies*dilution factor) / volume of culture plate.

The pathogenicity test on entire plants involved spraying a fungal suspension containing 1*10^6^ conidia/ml onto 75-day-old plants every alternate day, with photographs taken after 20 days. Additionally, for the light-dark fungal bioassay, 10-day-old seedlings grown in light and those exposed to 48 hours of darkness were infected with the fungus and subsequently utilized for biochemical analysis.

### Histochemical detection of superoxide and hydrogen peroxide accumulation

To detect the production of superoxide radicals, NBT staining was performed according to the method described in Jabs et al., 1996 with slight modifications. The leaves were immersed in 50 mM potassium phosphate buffer (pH 7.8) containing 0.1 mg ml^−1^ nitrobluetetrazolium (NBT) at 25°C in the dark for 24h. Stained samples were bleached in 80% ethanol and incubated at 70°C for 10–20 min. The accumulation of hydrogen peroxide (H_2_O_2_) in the samples was measured by using DAB staining. It was performed according to the method (Daudi and O’Brien, 2012). The leaves were immersed in 1 mg ml^−1^ DAB solution. To enhance the uptake of DAB, samples were vacuum infiltrated for 5–10 min. Staining was carried out in dark for 24h with mild shaking. Finally, the samples were de-stained using a 1:1:3 mixtures of acetic acid, glycerol and ethanol at 95°C for 15 min.

### Estimation of MDA content

Lipid peroxidation assay is a biochemical marker for oxidative stress, which is estimated by malondialdehyde (MDA) produced using the thiobarbituric acid (TBA) method (Baryla et al., 2000). Briefly, 0.1 g of sample was ground into powder using liquid nitrogen and was homogenized in 1 ml 0.1% (w/v) trichloroacetic acid (TCA). The homogenate was centrifuged at 10,000xg for 10 min. The supernatant was mixed with 4 ml of 20% (w/v) TCA containing 0.5% (w/v) TBA, heated in a boiling bath (95^0^C) for 15 min and then allowed to cool rapidly in an ice bath. The mixture was centrifuged at 10,000xg for 5 min and the resulting supernatant was used for the determination of MDA content. The concentration of MDA was calculated by measuring absorbance at 532 nm (correction was made by subtracting absorbance at 600 nm for turbidity) by using an extinction coefficient of 155 mM^−1^ cm.^−1^

### Cell wall degrading enzyme assay

Cellulase activity was measured by determining the liberated reducing end products using glucose as standards (Durbin and Lewis., 1988). The reaction mixture (0.5 ml) contained 1% substrate, 0.05 M sodium acetate buffer (pH 5.5) and a suitable amount of crude extract. Assays were carried out at 37°C for 1 h. Then 0.5 ml DNS (dinitrosalicylic acid) reagent was added to each tube. The tubes were heated in a boiling water bath for 10 min. After cooling to room temperature, the absorbance was measured at 560 nm. Substrates used was CM-cellulose for cellulase. One unit of enzyme activity was defined as the amount of enzyme which liberated 1 mol of reducing sugar per h under standard assay conditions.

### Total protein extraction and western blot analysis

Whole seedlings (300 mg) were frozen in liquid N_2_ and ground in extraction buffer (50 mM Tris (pH 7.5), 150 mM NaCl, 1 mM EDTA, 10% glycerol, 5 mM DTT, 1% protease inhibitor cocktail (Sigma-Aldrich), 0.1% Triton-X100). The homogenate was transferred to a fresh eppendorf tube and centrifuged at 20,000 g for 12 min at 4 °C. The supernatant was transferred to a new tube, and an aliquot (5 µl) was used to estimate the protein concentration using Bradford assay (Bradford., 1976). The mixture of Laemmli sample buffer (2X) and the protein sample were boiled at 96 °C for 5 min. The protein samples (10 µg) were separated on SDS–PAGE (12%) gel at 150 V for 2 h and transferred onto a PVDF membrane at 25 V for 2 h in transfer buffer (3.03g Tris–HCl, 14.4g glycine, 20% methanol) using a semi-dry transfer apparatus (Bio-Rad). After transfer, the membranes were incubated in the blocking solution (3% non-fat dry milk in Tris-buffered saline containing 0.05% Tween-20) for 1 h at room temperature. The blots were incubated in primary antibodies diluted in TBS-T (1:10000 (v/v) for anti-actin, 1:500 (v/v) for anti-HY5 overnight, followed by 5 min washing with wash buffer (Tris-buffered saline containing 0.05% Tween-20) thrice. The secondary antibody, conjugated with horseradish peroxidase (anti-rabbit IgG-HRP and anti-mouse IgG-HRP) diluted (1:10,000 times) in blocking buffer was incubated for 1 h at room temperature, followed by 5 min washing with wash buffer three times, followed by visualization using ChemiDoc (MP system, Bio-Rad) after incubating with luminol/enhancer and peroxide buffer (1:1 ratio). The western blot images were captured using Image Lab version 5.2.1 build 11 (Bio-Rad Laboratories). The commercial antibodies used in the analysis were Anti-Actin Antibody (A0480; Sigma-Aldrich), anti-HY5 (AS12 1867; Agrisera).

### Statistical analysis

Data are plotted as means±SE (n=3) in the figures. For CFU count and hypocotyls, root, rosette diameter, seed size n=5 and n=10 were taken into the consideration, respectively. Data of flavonol and SGA accumulation were evaluated for statistical significance using the Two-tailed Student’s t-test using Graphpad prism 8.05 software. Asterisks indicate significance levels with (* P < 0.1; ** P < 0.01; *** P < 0.001) as confidence intervals.

## Author contributions

P.K.T. designed and supervised this study. H.S. designed and performed most of the experiments; R.S.K., T.D., and D.S. provided help in conducting various experiments; P.K.T. analyzed the data; H.S. and P.K.T. wrote the article; all authors read, contributed, and approved the article.

## Acknowledgment

P.K.T. acknowledges CSIR and DBT, New Delhi, for financial support in the form of projects on pathway engineering and genome editing. P.K.T. also acknowledges Science and Engineering Research Board, New Delhi for JC Bose National Fellowship (JCB/2021/000036). H.S. acknowledges DBT, New Delhi for Senior and Junior Research Fellowship respectively. Authors also acknowledge Dr. Manju Singh and Dr. Ratnashekhar CH from CSIR-CIMAP for help in metabolite profiling.

## Supplemental Data

**Supplemental Figure S1.** Estimation of flavonol content in WT tomato.

**Supplemental Figure S2**. Regulation of flavonoid biosynthesis genes by light in AtMYB12 overexpression line of tomato.

**Supplemental Figure S3**. Flavonol content modulation by light in AtMYB12 overexpression line of tomato.

**Supplemental Figure S4.** Sequence analysis of *SlHY5* gene.

**Supplemental Figure S5.** Evolutionary analysis of SlHY5 by maximum likelihood method.

**Supplemental Figure S6**. Complementation of *Arabidopsis* and tobacco *hy5* mutant by SlHY5.

**Supplemental Figure S7**. Rossette diameter of complemented lines of *Arabidopsis*.

**Supplemental Figure S8**. The phenotype of mature *Arabidopsis*.

**Supplemental Figure S9**. Seed size in *Arabidopsis* is regulated by SlHY5.

**Supplemental Figure S10.** Schematic representation of flavonoid metabolism pathway for three different plants used in the study.

**Supplemental Figure S11**. Relative transcript level of flavonoid biosynthesis genes in complemented lines of *Arabidopsis* and tobacco.

**Supplemental Figure S12**. Accumulation of flavonoids in complemented lines of *Arabidopsis* and tobacco.

**Supplemental Figure S13.** High-performance liquid chromatography (HPLC) of *Arabidopsis* plants.

**Supplemental Figure S14**. Light-induced expression of flavonoid biosynthetic genes in *Arabidopsis* plants.

**Supplemental Figure S15**. Biochemical analysis of WT, *hy5*, and complemented lines.

**Supplemental Figure S16**. Position of gRNA in *SlHY5* sequences and amino acid sequence of the truncated protein in CRISPR-edited plants.

**Supplemental Figure S17**. Phenotype of transgenic tomato plants.

**Supplemental Figure S18**. SlHY5 modulates the gene expression of flavonoid biosynthesis genes.

**Supplemental Figure S19**. Differential accumulation of metabolite levels in transgenic tomato plants.

**Supplemental Figure S20**. Recombinant protein expression and purification of SlHY5.

**Supplemental Figure S21**. SlHY5 physically binds to *SlMYB12* promoter.

**Supplemental Figure S22**. SlHY5 physically binds to promoters of SGA genes.

**Supplemental Figure S23.** Fold enrichment of SlHY5 to *SlMYB12* promoter.

**Supplemental Figure S24.** SlHY5 coordinates with promoter of SGA and flavonoid biosynthesis genes.

**Supplemental Figure S25.** Light-responsiveness of flavonol pathway genes.

**Supplemental Figure S26.** SlHY5 modulates flavonoid content in tomato.

**Supplemental Figure S27.** Fungal bioassay of tomato transgenic lines.

**Supplemental Figure S28.** Differential ROS production in SlHY5OX and *SlHY5^CR^* plants after fungal infection.

**Supplemental Figure S29.** Cell wall degrading enzyme assay in SlHY5OX and *SlHY5^CR^* plants after fungal infection.

**Supplemental Figure S30.** Phenotypic analysis of SlHY5OX and *SlHY5^CR^* plants after fungal infection.

**Supplemental Figure S31.** HY5-induced light-dependent SGA and flavonoid biosynthesis as a fungal tolerance mechanism.

**Supplemental Table S1**. Putative cis-acting light-responsive elements in *SlGAME4* promoter.

**Supplemental Table S2**. Putative cis-acting light-responsive elements in *SlGAME11* promoter.

**Supplemental Table S3**. Putative cis-acting light-responsive elements in *SlGAME12* promoter.

**Supplemental Table S4**. Putative cis-acting light-responsive elements in *SlGAME17* promoter.

**Supplemental Table S5**. Putative cis-acting light-responsive elements in *SlC5SD* promoter.

**Supplemental Table S6**. Putative cis-acting light-responsive elements in *SlGAME9* promoter.

**Supplemental Table S7**. Putative cis-acting light-responsive elements in *SlMYB12* promoter.

**Supplemental Table S8.** Oligonucleotides used for development of constructs and expression analysis.

## Data availability

All data generated or analyzed during this study are included in this published article (and its supplementary information files).

## Conflict of interest

The authors declare that they have no conflict of interest.

